# History-dependent changes of Gamma distribution in multistable perception

**DOI:** 10.1101/2020.08.06.239285

**Authors:** Alexander Pastukhov, Malin Styrnal, Claus-Christian Carbon

## Abstract

Multistable perception – spontaneous switches of perception when viewing a stimulus compatible with several distinct interpretations – is often characterized by the distribution of durations of individual dominance phases. For continuous viewing conditions, these distributions look remarkably similar for various multistable displays and are typically described using Gamma distribution. Moreover, durations of individual dominance phases show a subtle but consistent dependence on prior perceptual experience with longer dominance phases tending to increase the duration of the following ones, whereas the shorter dominance leads to similarly shorter durations. One way to generate similar switching behavior in a model is by using a combination of cross-inhibition, self-adaptation, and neural noise with multiple useful models being built on this principle. Here, we take a closer look at the history-dependent changes in the distribution of durations of dominance phases. Specifically, we used Gamma distribution and allowed both its parameters – shape and scale – to be linearly dependent on the prior perceptual experience at two timescales. We fit a hierarchical Bayesian model to five datasets that included binocular rivalry, Necker cube, and kinetic-depth effects displays, as well as data on binocular rivalry in children and on binocular rivalry with modulated contrast. For all datasets, we found a consistent change of the distribution shape with higher levels of perceptual history, which can be viewed as a proxy for perceptual adaptation, leading to a more normal-like shape of the Gamma distribution. When comparing real observers to matched simulated dominance phases generated by a spiking neural model of bistability, we found that although it matched the positive history-dependent shift in the shape parameter, it also predicted a negative change of scale parameter that did not match empirical data. We argue that our novel analysis method, the implementation is available freely at the online repository, provides additional constraints for computational models of multistability.

**Author Summary:** Multistable perception occurs when one continuously views a figure that can be seen in two distinct ways. A classic old-young woman painting, a face-vase figure, or a Necker cube are examples easy to find online. The endless spontaneous switches of your perception between the alternatives inform us about the interplay of various forces that shape it. One way to characterize these switches is by looking at their timing: How long was a particular image dominant, how did that reflect what you have seen previously, the focus of your attention, or the version of the figure that we showed you? This knowledge allows us to build models of perception and test them against the data we collected. As models grow more elaborate, we need to make tests more elaborate as well and for this, we require more precise and specific ways to characterize your perception. Here, we demonstrate how your recent perceptual experience – which of the alternative images did you see and for how long – predicts subtle but consistent changes in the shape of the distribution that describes perceptual switching. We believe it to be a more stringent test by demonstrating how a classical model of bistability fails on it.

## Introduction

We rely on unambiguous perception to be able to initiate actions in a very fast and clearly directed way. The real challenge of perception is to generate such a clear and single representation of the outside world that is based on inherently noisy and ambiguous sensory information. As the mental act of creating such perception must evidently be based on perceptual models, the representation will not always match the actual state of the external world leading to characteristic and highly informative visual illusions [1]. Sometimes, sensory information is compatible with several comparably likely perceptual interpretations [2]. A few of these so-called multistable figures can be seen in **Figure 1A-C**, and many more stimuli do exist and are systematically investigated, including an auditory [3], tactile [4], and even olfactory bistable stimuli [5]. Thus, multistable perception is particularly interesting because it is a general phenomenon, so understanding it advances our knowledge about the architecture of our perceptual system and perceptual decision making.

**Figure 1.**
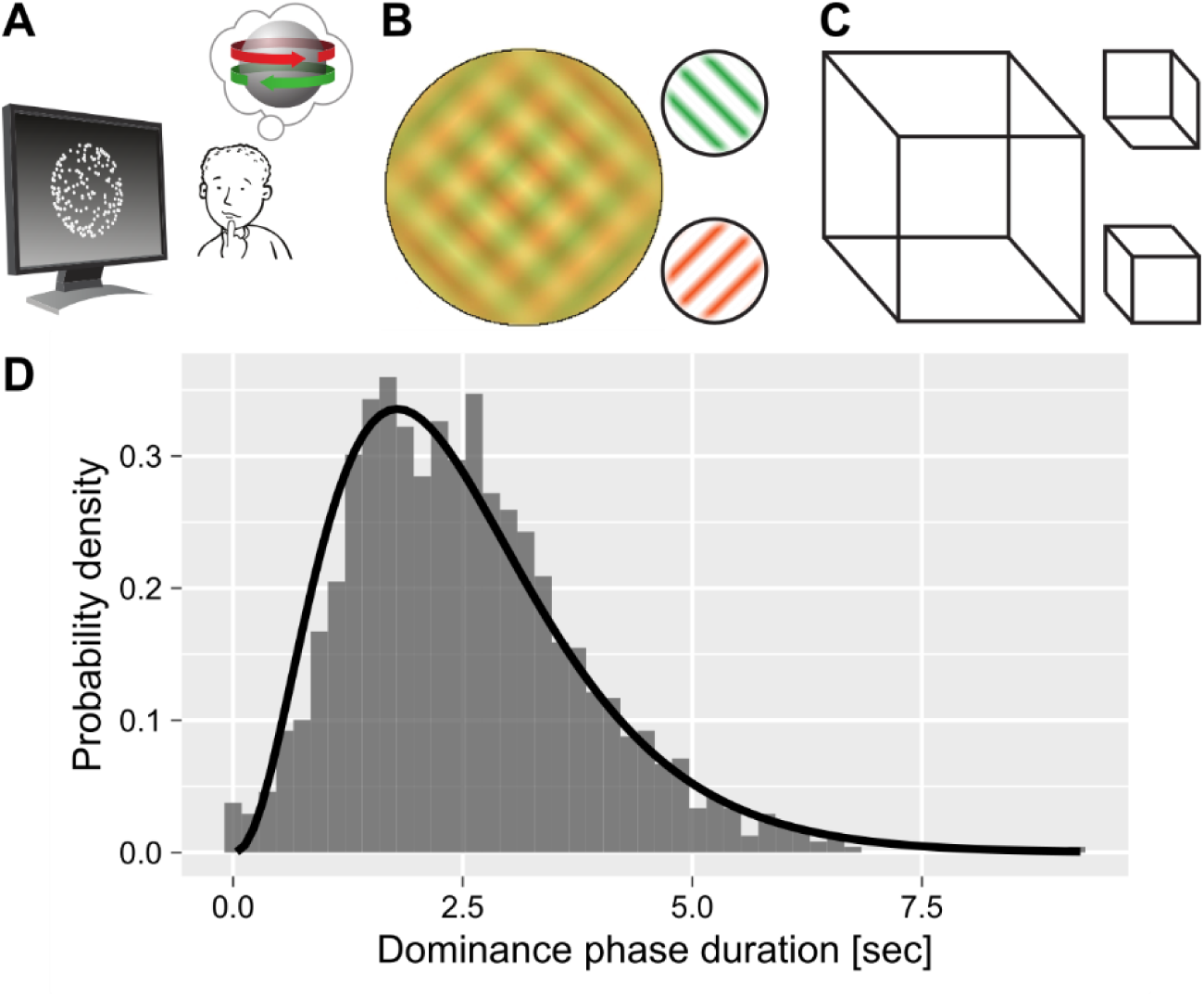
Bistable displays (A, B, C) and a typical distribution of dominance phase durations (D). A) In the kinetic-depth effect, a 2D onscreen-motion produces an alternating perception of a rotating object. B) Binocular rivalry produces alternating dominance of two orthogonal gratings. C) Necker cube can be perceived in two orientations. D) Distribution of dominance phases for a participant, who viewed kinetic-depth effect displays, with an overlaid fitted gamma distribution.

When multistable displays are presented continuously, their perception is often characterized by a distribution of duration times or, conversely, by the alternation rate. Although duration times vary strongly between subjects, the shape of the distributions is remarkably similar [6], see **Figure 1D**. This consistency is viewed as a “hallmark” of multistable perception [7] and is frequently used as a first check when characterizing a new multistable display [8]. There is a debate about which theoretical distribution fits the empirical dominance phase durations the best with many suggestions that included gamma, exponential, Weibull, normal, Capocelli-Ricciardi, beta rate, ex-gaussian, and log-normal distributions [9–15]. Among those, the gamma distribution is generally considered to be the canonical distribution for fitting the data [9,16,17]. Its specific appeal is that it allows one to conceptualize perceptual alternations as a Poisson process that occurs after α discrete stochastic events, which occurr independently [9,17]. Interestingly, this conceptualization predicts that the shape parameter must be an integer number but the evidence for this is not conclusive [16].

Another important property of dominance phase durtions is a subtle but consistent dependence of their duration on prior perceptual history. This dependence is particularly evident when an unambiguous display precedes the bistable one. Here, an artificially prolonged dominance of a particular percept leads to longer dominance of the alternative percept during a consequent continuous presentation [18,19]. Although less pronounced, the same serial dependence is observed when fully ambiguous bistable displays are viewed continuously. A typical approach is to compute an autocorrelation of dominance phase durations with different lags [17] and a typical result is a small (0.1-0.2) but significant and consistently positive autocorrelation for lag 1 [20]. In other words, longer dominance phases tend to be followed by similarly long ones and, conversely, short phases tend to be followed by comparably short ones. Alternatively, prior perceptual history can be quantified using a leaky integrator with a fitted time constant [21]. The latter approach tends to produce a higher correlation between preceding history and the following dominance phase. Thus, it is likely to be a more accurate estimate of the perceptual history and, therefore, was used in the current study.

Typical models of bistable perception assume that it is borne out as an interplay between cross-inhibition, self-adaptation, and noise [22–25]. The latter appears to be the main driver of perceptual alternations [26], whereas the former ensures perceptual exclusivity. The dependence of dominance phase duration’s on prior perceptual history described above is thought to reflect noisy self-adaptation, so that high levels of adaptation of a particular percept allow for prolonged dominance of the alternative [20]. Conversely, similarly low levels of adaptation minimize its effect, so that perceptual alterations are driven primarily by stochastic factors [21,27].

As summarized above, the prior work showed that the mean of distribution of dominance durations is consistently shifted due to changes in adaptation levels. However, for the gamma distribution, a particular change of the mean is consistent with various combinations of changes to its shape and scale parameters (**Figure 2**). For example, a higher contrast in binocular rivalry displays results in shorter mean dominance phase durations, which is a product of a simultaneous increase in the shape and decrease in the scale parameters [20]. This change is more evident when the balance between adaptation and noise is manipulated by reducing the former, leading to an exponential distribution, *e*.*g*., gamma distribution with shape=1 [27]. Similarly, modelling demonstrates a consistent change in the gamma distribution when the balance between noise and adaptation is manipulated ranging from exponential, when dominated by noise, to normal, when dominated by adaptation [25]. Thus, the parameters of the gamma distribution should also reflect the changing levels of adaptation and, therefore, the shift in the balance between adaptation and noise.

**Figure 2.**
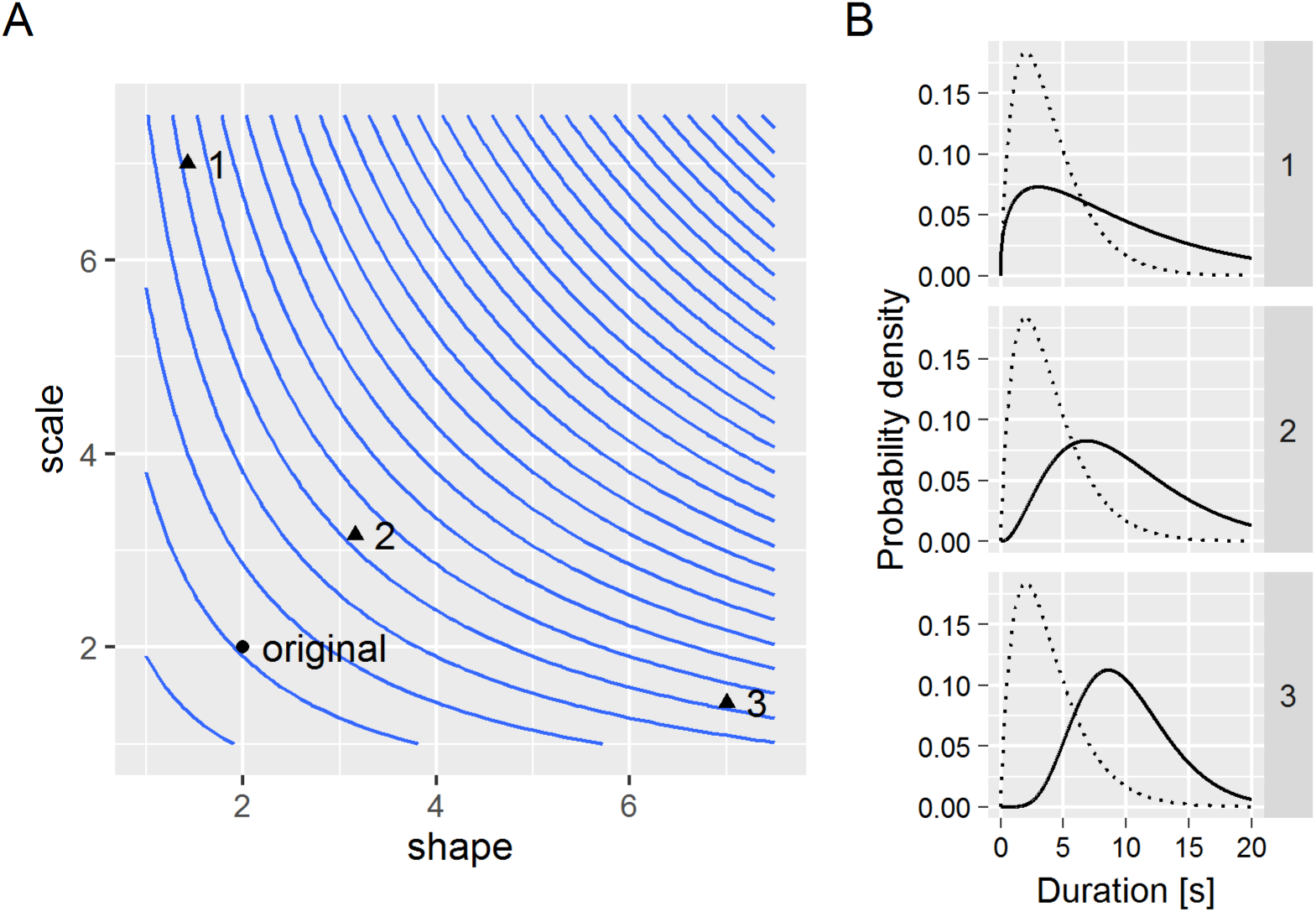
The same change in the mean of the gamma distribution can be implemented as various combinations of changes to its shape and scale parameters. A) Isolines indicate parameters’ combinations that produce the same mean. B) Three example distributions that have the same higher mean as compared to the original (dashed line).

In the present study, we sought to characterize the changes to gamma distribution using prior perceptual history computed via leaky integrator [21]. Specifically, we computed perceptual history for two timescales – short (τ=0.9 <T_dom_>, where <T_dom_> stands for mean dominance duration) and long (τ=30 <T_dom_>). Then, we used a hierarchical Bayesian model with gamma distribution allowing both parameters to vary with respect to perceptual history, stimulus contrast, and age of participants. We demonstrate that higher levels of perceptual history are associated with positive changes in shape parameter, leading to more “Gaussian-like” distribution, but we found no consistent changes in scale parameter. Finally, we fitted the same model on data simulated via a spiking neural model of bistability [22]. We observed that these did not fully match empirical observations, as positive changes in shape were accompanied by equally strong, and absent for empirical data, negative changes in the scale parameter. We conclude that our novel method improves our ability to characterize time-series of perceptual alternations and provides additional constraints for their models.

## Materials and Methods

### Data sets

We used two previously published data sets together with newly measured data (see **Table 1** and the description below). The first data set, reported in [28], contains results for binocular rivalry (BR), kinetic-depth effect (KD), and Necker cube (NC) displays measured in an adult population. Second, reported in [29] used BR display measured in three age groups (9, 12, and 21 years old). For details please refer to the corresponding papers. We have included [28] and the new data set in the open data repository. However, the data set that contained children was not included as we had no permission to disclose it.

**Table 1.**
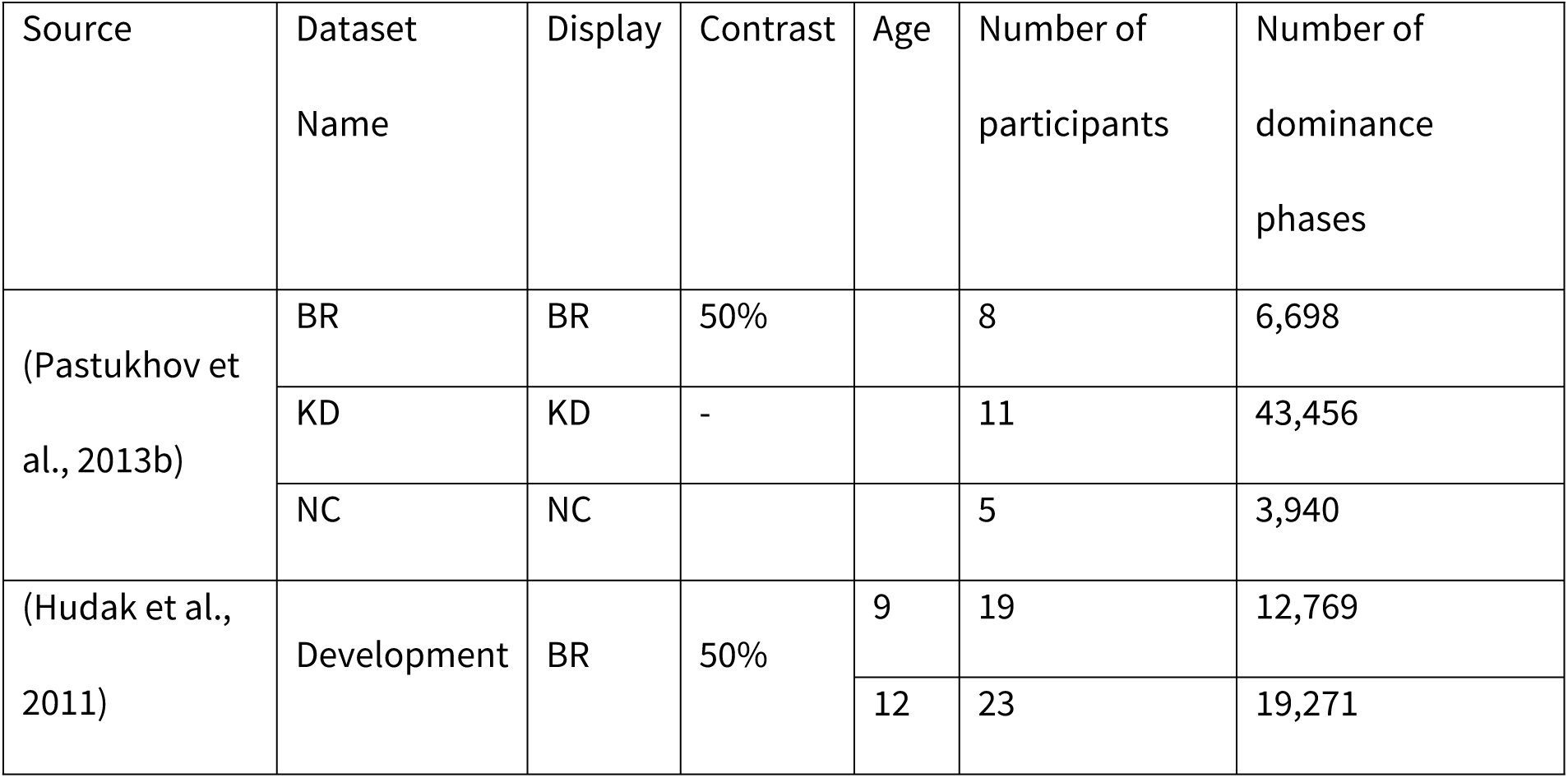
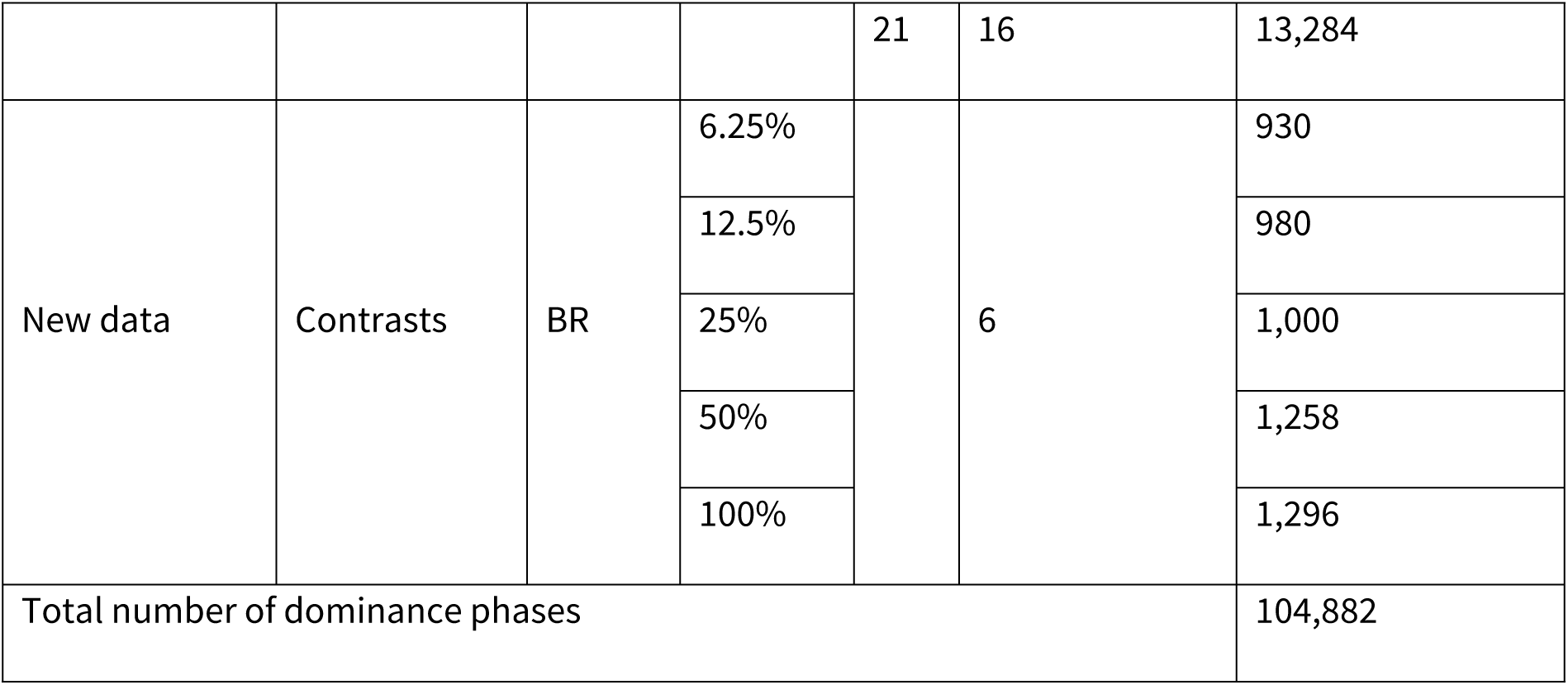
Summary of data sets used in the study. BR – binocular rivalry, KD – kinetic depth effect display, NC – Necker cube.

For the new data set, we used binocular rivalry display with a procedure similar to that used in [28] but with five contrasts levels (6.25%, 12.5%, 25%, 50%, 100%). Binocular rivalry stimulus consisted of two orthogonally oriented gratings (radius 0.9°, spatial frequency 2 cycles/°, one was tilted by 45° clockwise and one 45° counter-clockwise). Participants viewed the display through a custom-made mirror stereoscope (75 cm eye-screen distance) and reported on the eye dominance using a keyboard, continuously pressing *left* when the counter-clockwise oriented grating was dominant, continuously pressing *right* when the clockwise-tilted grating was dominant. Lack of key presses indicated mixed phases. Each presentation lasted for two minutes. An experimental session consisted of ten blocks, so that each contrast condition was repeated twice. The order was randomized, so that the five contrast conditions were shown in the random order and then again in the reverse order. The data was collected at Magdeburg University.

Six participants took part in the experiment. All participants signed the informed consent prior to the experimental session and received monetary compensation. They had normal or corrected-to-normal vision. All procedures were in accordance with the national ethical standards on human experimentation and with the Declaration of Helsinki of 1975, as revised in 2008, and were approved by the medical ethics board of the Otto-von-Guericke Universität, Magdeburg: “Ethikkomission der Otto-von-Guericke-Universität an der Medizinischen Fakultät.”

### History computation

The perceptual history that preceded each dominance phase duration was computed via leaky integrator with an exponential kernel as in [21]. The history was computed for two timescales, with the short timescale history τ=0.9·<T_dom_> and the long timescale τ=30·<T_dom_>, where <T_dom_> is the mean dominance phase duration for a given participant, display, and contrast. We selected the short timescale time constant based on the results of [21] and so as to represent the most recent time history. Conversely, the purpose of the long timescale was to capture a possible slow drift in dominance phase durations [30]. Once the perceptual history was computed, we excluded the first two clear percepts of each block from the analysis as they did not have any preceding history for that state.

### Simulated data

We have generated simulated data using a custom implementation of a spiking neural model of bistability [22] based on the code provided by Stepan Aleshin and Jochen Braun from Cognitive Biology group at Magdeburg University. The model was fitted individually for each participant and display from the first data set (Pastukhov et al., 2013b). We used a genetic algorithm [31] with the Kolmogorov-Smirnov test as a fitness function, to match the dominance phase distributions for the experimental and the simulated data as closely as possible. Note, however, that the fitness function did not include any information on the history dependence. The population size was 50 and the number of iterations 100. After the final iteration, we used best model parameters to generate a time series with 1100 clear percepts (first 100 percepts were discarded to account for the initial “burn-in” period of the model), see **Figure 3**. Because the simulated data is stationary and does not exhibit any long-timescale drift, we computed perceptual history only for the short timescale (τ=0.9·<T_dom_>).

**Figure 3.**
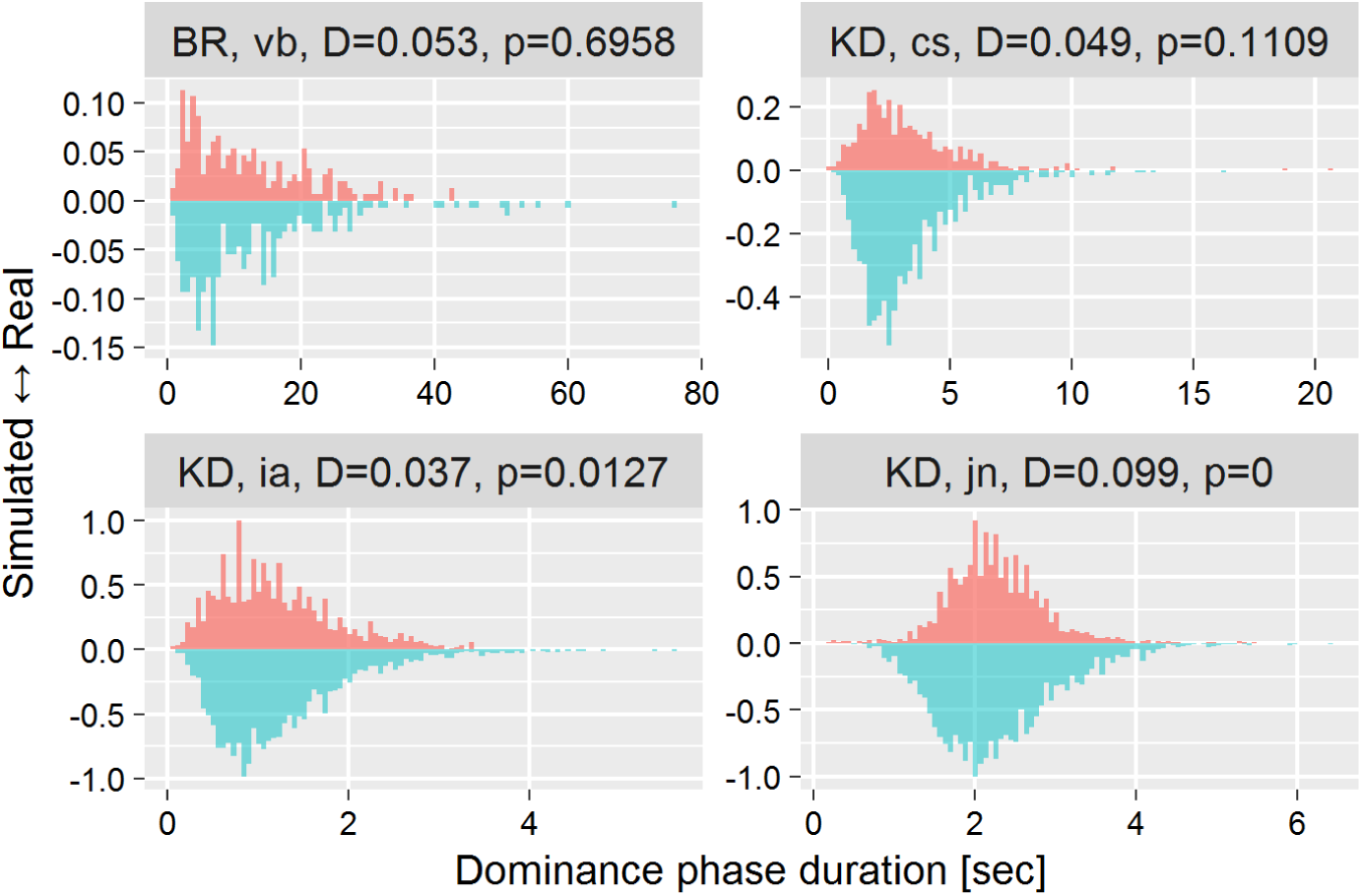
Example distributions of dominance phase durations for the experimental (upwards) and matching simulated (downwards) data for four participants. We picked four participants based on the Kolmogorov-Smirnov test p-value from worst-matched (BR, vb) to best-matched (KD, jn) with two intermediate cases (at 1/3 and 2/3). The title above each plot shows the display, participant code, Kolmogorov-Smirnov test statistics and the p-value.

### Model

We fitted each data set separately using a Bayesian hierarchical generalized linear model. Models for two parameters of gamma distribution – shape (*k*) and scale (Θ) – were identical. For all groups, each parameter depended on prior perceptual history at two timescales, except for the simulated data, there they depended only on short-term history (see above). For *contrasts* and *development* data we also included the main effects of, correspondingly, contrast and age.

Before fitting the model, the data were centered as follows. Prior history was centered at its median value for each participant × timescale. For *contrasts* group, the contrast was centered at 50% value. For the *age* group, age was centered at a proposed mean young adult age of 21 years old.

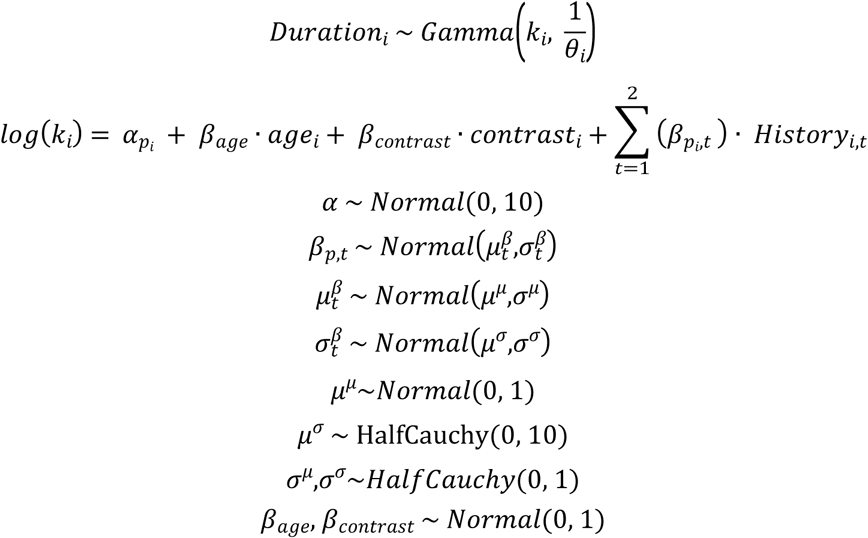

where *i* is the row index, *t* is the time scale index (1..2 range), *k*_*i*_ is the shape parameter for the *i*^*th*^ row, *p*_i_ is the participant for the *i*^*th*^ row, *age*_*i*_ is participant’s age, *contrast*_*i*_ is the display contrast, and *History*_*i,t*_ is the computed history value for *i*^*th*^ row and *t*^*th*^ time scale. The model for the scale parameter (Θ_i_) was identical.

The model was programmed and sampled using Stan probabilistic programming language [32].

### Software

The analysis and modelling were performed in R 4.0.0 [33] using *Tidyverse* collection of packages [34]. The model was programmed and sampled using *rstan* [35]. The spiking neural model of bistability was implemented using *Rcpp* [36]. We used *GA* package [37] for genetic algorithm optimization.

### Open Practices Statement

Two out of three data sets (excluding the developmental data), the analysis code and the model code are available under Creative Commons Attribution 4.0 International Public License at https://osf.io/js3wv or https://github.com/alexander-pastukhov/history-dependent-gamma.

## Results

Our aim was to investigate how perceptual history affects the distribution of the following dominance phase durations. Specifically, we assumed that this distribution is well-approximated by Gamma (see Introduction for rationale) and we were interested in the history dependence of the two individual parameters, shape (k) and scale (θ). To this end, for each dominance phase, we computed perceptual history for both short and long timescales. Then, we fitted each dataset using a hierarchical Bayesian model where both parameters linearly depended on both timescales. In addition, we included the effect of contrast and participants’ age for, respectively, *contrast* and *development* data sets.

The shape and scale parameters of the fitted gamma distributions (without effects of history, contrast, and age) for all participants are plotted in **Figure 4A**. They show typical values with most shape parameters clustering between 2 and 4 and similarly tight clustering for the scale parameter. For the data set that used *contrast* manipulation, the effect of contrast matches that reported by van Ee [20]. Specifically, higher contrast led to a positive change for the shape parameter but a negative one for scale parameter (**Figure 4B**, left panel). However, we found no effect of age on either parameter of the gamma distribution (**Figure 4B**, right panel).

**Figure 4.**
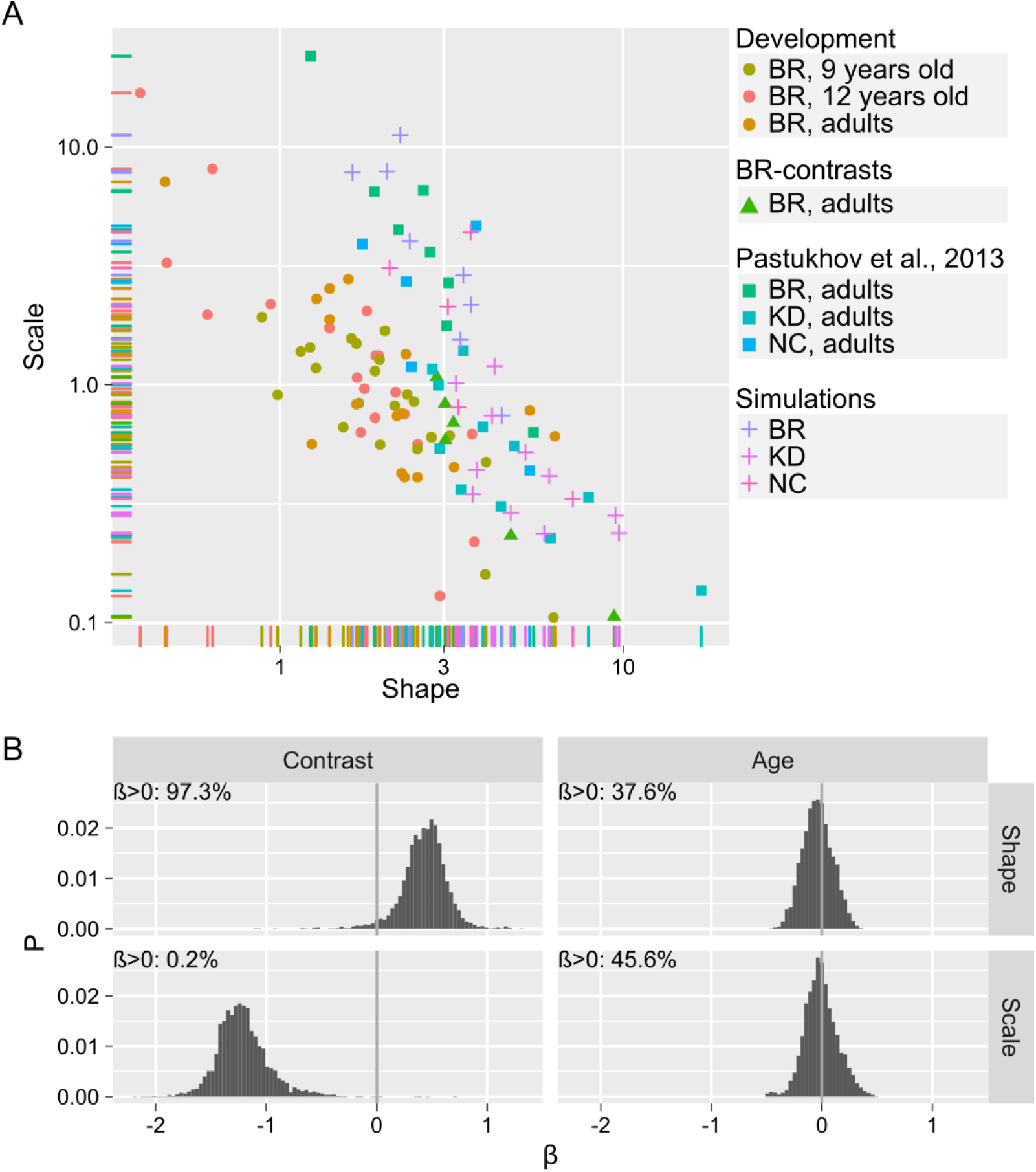
Intercepts-only distribution parameters and main effects of contrast and age. A) The scatterplot shows the intercepts-only parameters of the fitted gamma distribution for each participant, including the simulated data. Shape and scale correspond to α^shape^ and α^shape^ except for *development* group data where it is equal to α^shape^ + β_age_· Age and α^scale^ + β_age_ · Age (see the model for further details). B) Posterior distributions for main effects of contrast (*Contrasts* data set) and age (*Development* data set) for shape and scale parameters of the gamma distribution. Text inset in the top left corner shows the percentage of β-values above zero.

The history dependence of the parameters for both timescales is summarized in **Figure 5**. For the short timescale perceptual history, we found a consistent positive shift for the shape parameter across all five data sets, although the effect was weaker for Necker Cube (NC). For the scale parameter, we also found a generally positive effect, although it was weaker and less consistent across the data sets than that for the shape. The effect of the history for longer time scale was more variable with mostly negative changes to both parameters indicating an overall speed-up trend for perceptual alternations over time.

**Figure 5.**
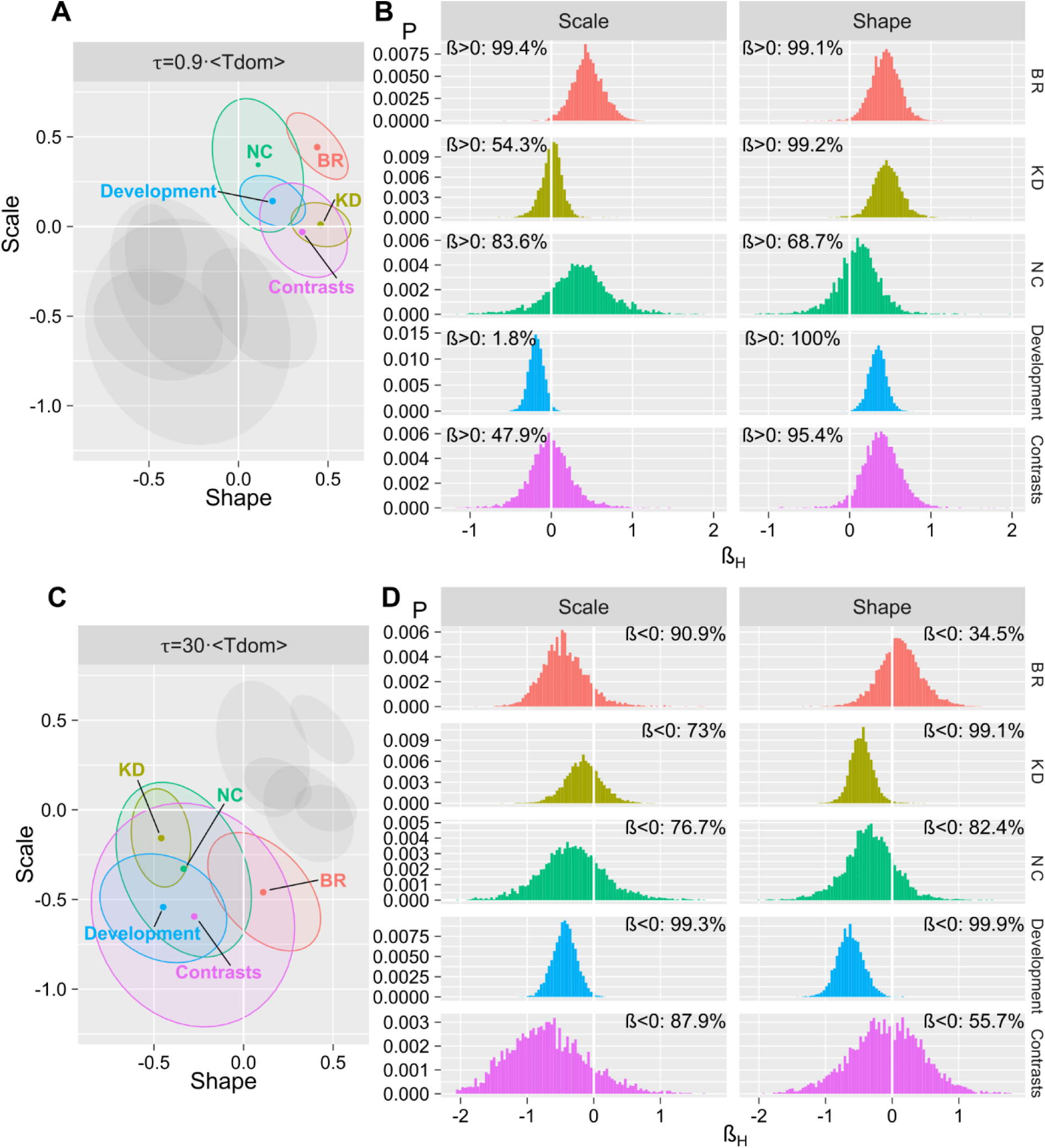
Posterior distributions for the main effect of prior history. A, B) short timescale (τ=0.9<T_dom_>), C, D) long timescale (τ=30<T_dom_>). A, C) Ellipses represent a fitted multivariate t-distribution at a level of 0.5. Gray ellipses depict results for the complementary timescale. B, D) Text insets show the percentage of β-values above or below 0.

We also compared the dominance phases of real observers with that generated by a spiking neural model of bistability [22]. The model was fit so as to match distributions of individual participants in three data sets (BR, KD, and NC) as closely as possible (see Methods for details). However, the simulated data was not optimized with respect to history-dependence, as the latter was not included in the fitness function. In addition, the selected model does not generate a long-term trend and, therefore, we opted to use only the short timescale history. The comparison of the main effect of perceptual history is presented in **Figure 6** with the results matching those of the real observers only partially. Although we found a similar positive shift for the shape parameter, we also observed a consistent *negative* history-dependent shift for the scale parameter. This contrasts simulated data with that from human observers, as for the latter we found a small and inconsistent but mostly positive change for the scale parameter (gray histograms in (**Figure 6**). Interestingly, the effect of short timescale history for the model was similar to that of contrast (**Figure 4B**).

**Figure 6.**
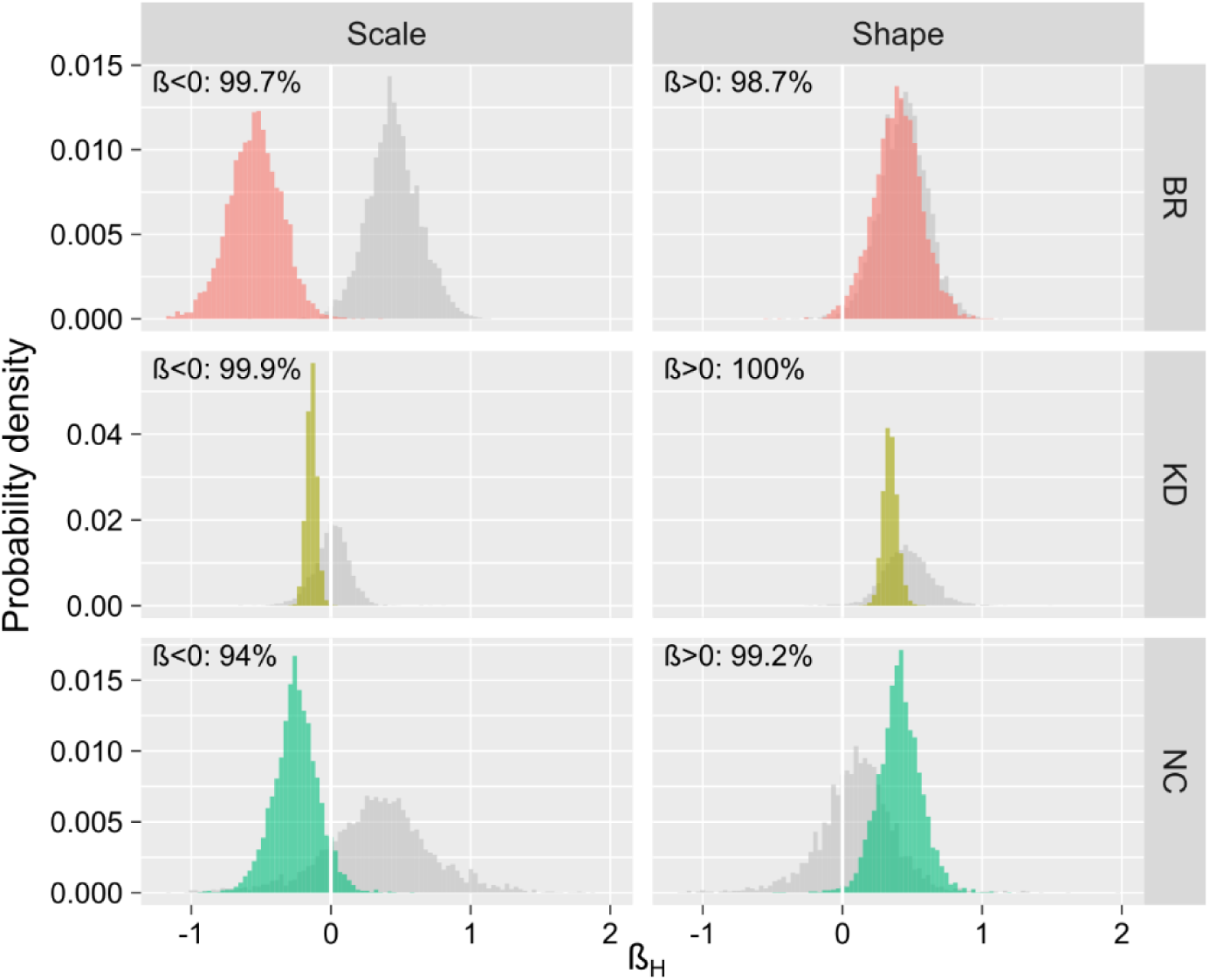
Simulated data, posterior distributions for short timescale history (τ=0.9<T_dom_>). Text inset in the top left corner shows the percentage of β-values above or below zero. The gray color shows posterior distributions for matching parameters and displays for the real participants (see Figure 5).

## Discussion

The aim of this study was to characterize how prior perceptual history affects the distribution of the following dominance phase durations. To this end, we fitted the distributions of dominance phase durations using gamma distribution with its parameters being linearly dependent on prior perceptual history at two timescales, as well as contrast and age for two data sets. The two timescales were short (τ=0.9<T_dom_>, where <T_dom_> is an average dominance phase duration) and long (τ=30<T_dom_>), so as to capture both effect of the immediately preceding dominance state and the overall trend. For the former, we observed a consistent positive dependence for the shape parameter and a weaker, less consistent but also positive dependence for the scale parameter. For the latter, both parameters had a negative dependence on the longer-term prior history, indicating a general trend for faster perceptual alternations.

Of the two timescales, we focused on in the present study, the short one is often presumed to reflect the effect of perceptual adaptation [20,21]. As we explained in the introduction, perceptual adaptation is considered to be one of the two major drivers of perceptual alternations alongside the neural noise. This conceptualization predicts that predictability of the individual dominance phase duration depends on the balance between the two. When one of the competing populations is strongly adapted, the dominance duration tends to be longer reflecting the greater influence of adaptation. Conversely, when adaptation levels of two competing representations are comparable, the perceptual alternations and, therefore, dominance phase durations are determined primarily by noise [21,26]. Our results appear to bear out this prediction, as the higher levels of prior perceptual history – a proxy for perceptual adaptation – are associated with consistent positive changes for the shape parameters but only weaker and less consistent ones for the scale parameter of the gamma distribution. The overall effect is that the distribution is more normal-like and, therefore, is more regular when history/adaptation is high [25]. Conversely, it is more exponential-like when history/adaptation is low and perceptual switches are driven primarily by noise [27]. In short, our results tested and validated prior assumptions about the roles of noise and adaptation in multistable perception and provide further means of characterizing it.

The proposed analysis also provides a useful benchmark for models of multistability. Here, we used a spiking neural model of bistability [22] that is capable of capturing various properties of dominance phase distribution for continuous viewing [25]. Multiple more capable models are built on the same idea of mixing self-adaptation, cross-inhibition, and neural noise [38–40]. There exist a plethora of alternative approaches including stochastic accumulation in multiple assemblies [6], Bayesian sampling [41,42], and even quantum theoretical network [43]. Our observations provide a more subtle characterization of the time-series and, therefore, a more sensitive test for all these models. Importantly, this is the test that the model by Laing and Chow appears to fail. Although the model does exhibit the positive history-dependence of the shape parameter, it also shows a strong and consistent *negative* history-dependence for the scale parameter. These results do not completely match those of human participants, indicating that the model is incapable of fully capturing the underlying dynamics. We must note, however, that the model parameters were not optimized for history-dependence only for the overall match of the distribution shapes.

Accordingly, the observed history-dependence in simulated data was not there by choice but was a product of the model itself. Therefore, it is possible that more thorough parameter-matching could produce a better match. Nonetheless, this discrepancy shows the usefulness of our approach for in-depth analysis of computational models of multistable perception.

To conclude, we presented a novel analysis method, the implementation is available freely at the online repository, which improves our ability to characterize the history-dependence of time-series of perceptual alternations, provides additional constraints for computational models of multistability.

## Acknowledgments

We thank Stepan Aleshin and Jochen Braun for providing us the code for the spiking neural model of bistability. I “thinking boy” image in Figure 1A is used under Public Domain license.

